# A drug repurposing screen identifies hepatitis C antivirals as inhibitors of the SARS-CoV-2 main protease

**DOI:** 10.1101/2020.07.10.197889

**Authors:** Jeremy D. Baker, Rikki L. Uhrich, Gerald C. Kraemer, Jason E. Love, Brian C. Kraemer

## Abstract

The SARS coronavirus type 2 (SARS-CoV-2) emerged in late 2019 as a zoonotic virus highly transmissible between humans that has caused the COVID-19 pandemic ^1,2^. This pandemic has the potential to disrupt healthcare globally and has already caused high levels of mortality, especially amongst the elderly. The overall case fatality rate for COVID-19 is estimated to be ∼2.3% overall ^3^ and 32.3% in hospitalized patients age 70-79 years ^4^. Therapeutic options for treating the underlying viremia in COVID-19 are presently limited by a lack of effective SARS-CoV-2 antiviral drugs, although steroidal anti-inflammatory treatment can be helpful. A variety of potential antiviral targets for SARS-CoV-2 have been considered including the spike protein and replicase. Based upon previous successful antiviral drug development for HIV-1 and hepatitis C, the SARS-CoV-2 main protease (Mpro) appears an attractive target for drug development. Here we show the existing pharmacopeia contains many drugs with potential for therapeutic repurposing as selective and potent inhibitors of SARS-CoV-2 Mpro. We screened a collection of ∼6,070 drugs with a previous history of use in humans for compounds that inhibit the activity of Mpro *in vitro*. In our primary screen we found ∼50 compounds with activity against Mpro (overall hit rate <0.75%). Subsequent dose validation studies demonstrated 8 dose responsive hits with an IC50 ≤ 50 μM. Hits from our screen are enriched with hepatitis C NS3/4A protease targeting drugs including Boceprevir (IC50=0.95 μM), Ciluprevir (20.77μM). Narlaprevir (IC50=1.10μM), and Telaprevir (15.25μM). These results demonstrate that some existing approved drugs can inhibit SARS-CoV-2 Mpro and that screen saturation of all approved drugs is both feasible and warranted. Taken together this work suggests previous large-scale commercial drug development initiatives targeting hepatitis C NS3/4A viral protease should be revisited because some previous lead compounds may be more potent against SARS-CoV-2 Mpro than Boceprevir and suitable for rapid repurposing.

## Introduction

The SARS virus and SARS-CoV-2, the cause of the COVID-19 pandemic, are zoonotic coronaviruses found in bats that can infect humans. Initial symptoms of SARS-CoV-2 infection include fever, myalgia, cough, and headache. Infection usually resolves without active medical intervention, but for a subset of cases infection can progress to viral pneumonia and a variety of complications including acute lung damage leading to death ^5^. While complications are atypical in most cases, mortality rates increase dramatically with the age and impaired health of infected patients. To date, much of our knowledge of COVID-19 virology has been inferred from the study of similar Severe Acute Respiratory Syndrome (SARS) coronavirus and related coronaviruses including Middle East Respiratory Syndrome [reviewed in ^6^]. Like all coronaviruses, SARS-CoV-2 exhibits an enveloped ribonucleoprotein helical capsid containing a single positive-stranded genomic RNA. Infection starts with receptor-mediated virus internalization, uncoating, and translation of the viral genome ^7^. Polyprotein cleavage by viral proteases yields a complement of viral structural and accessory proteins. This polyprotein cleavage is mediated by the main viral protease (Mpro), a chymotrypsin-like protease responsible for endoproteolytic cleavages of viral polyproteins producing functional viral proteins. Recent structural biology work has solved the crystal structure of SARS-CoV-2 Mpro yielding structural insights into Mpro function ^8,9^.

Antiviral drugs effective for COVID-19 would have a broad impact on global healthcare in the current coronavirus pandemic. Existing antiviral drugs on the market target a wide variety of both RNA and DNA viruses. Probably the most successful approved antivirals are protease inhibitors such as atazanavir for HIV-1 and simeprevir for hepatitis C. [reviewed in ^10^ and ^11^]. Other conceptual COVID-19 antiviral targets include the host ACE2 receptor, viral replicase, and viral genome encapsidation. However, previous work with other RNA viruses suggest that Mpro function is essential for viral replication and readily targetable using existing technology. Thus, while there are many potentially targetable activities for COVID-19, the coronavirus Mpro seems a likely choice for rapid drug development.

To accelerate drug development we employed a drug repurposing strategy, an approach of utilizing previously approved drugs for new indications ^12,13^. Previous work suggests libraries enriched with known bioactive drug-like compounds provide the best opportunity for finding new lead compounds ^14,15^. Thus we attempted the selective optimization of side activities (SOSA) approach ^16^ as a rapid and cost effective means to identify candidate hits while minimizing the number of compounds screened. The SOSA approach proceeds by two steps. First a limited set of carefully chosen, structurally diverse, well-characterized drug molecules are screened; as approved drugs, their bioavailability, toxicity and efficacy in human therapy has already been demonstrated ^16,17^. To screen as much of the available approved drug space as possible in an easily accessible format we chose to screen the Broad Institute Drug Repurposing Library (6070 compounds, see **Table S1**) ^18^. This represents about half of the approximately 14,000 approved or experimental drugs known to human clinical medicine ^19^. There are significant cost and time advantages realized by drug repurposing as it can accelerate the preclinical phase of development and streamline clinical trials to focus on efficacy rather than safety.

Repositioning existing approved drugs with the capacity to inhibit COVID-19 virus replication and infection would be of profound utility and immediately impact health care in the current pandemic. There are no drugs in clinical use specifically targeting coronavirus replication. The major advantage of the approach taken here is that by screening drugs with a history of previous clinical use, we will be focusing on compounds with known properties in terms of pharmacokinetics (PK), pharmacodynamics (PD) and toxicity. Thus, the Broad Repurposing Library we screened consists of compounds suitable for rapid translation to human efficacy trials.

## Results

### Development of fluorescent Mpro Assays

We began assay development by selecting potentially suitable synthetic Mpro substrates and compared catalyzed hydrolysis curves between 5 fluorescently labeled substrates (Ac-Abu-Tle-Leu-Gln-AFC ^20^, DABCYL-VKLQ-EDANS, Ac-VKLQ-AFC, DABCYL-TSAVLQSGFRKM-EDANS^21^, and MCA-AVLQSGFR-K(Dnp)-K-NH2)^22^. We chose to use the recently published Ac-Abu-Tle-Leu-Gln-AFC (Abu=2-Aminobutyrate, Tle=tButylglycine) synthetic non-canonical amino-acid containing peptide as Mpro more readily cleaves this preferred sequence as compared to the native VKLQ sequence ^20^(Fig 1A). Substrates DABCYL-TSAVLQSGFRKM-EDANS and MCA-AVLQSGFR-K(DnP)-K-NH2 had drastically lower rates of Mpro catalyzed hydrolysis and were not considered further in our assay development (Fig 1A). To determine concentration ratios between Mpro and substrate, we next preformed a two-dimensional titration and chose 625nM Mpro and 8µM substrate for a balance of relatively modest Mpro protein requirement and a robust fluorescence intensity (Fig 2B). Before screening the Broad library, we piloted our assay conditions against the NIH Clinical collections library (∼650 compounds) and calculated our Z’-factor for each plate at 0.780 and 0.784 (Fig 1C and D). Z’-factor is a score of suitability of assays for high-throughput screening and is derived from the equation 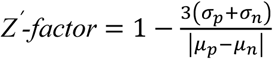, where σ = standard deviation, µ=mean, p=positive controls, and n=negative controls. A score greater than 0.5 indicates a screenable assay. Although no promising compounds were identified from this smaller library, it demonstrated that our assay was sufficiently robust for screening the much larger Broad Repurposing library.

**Fig 1.**
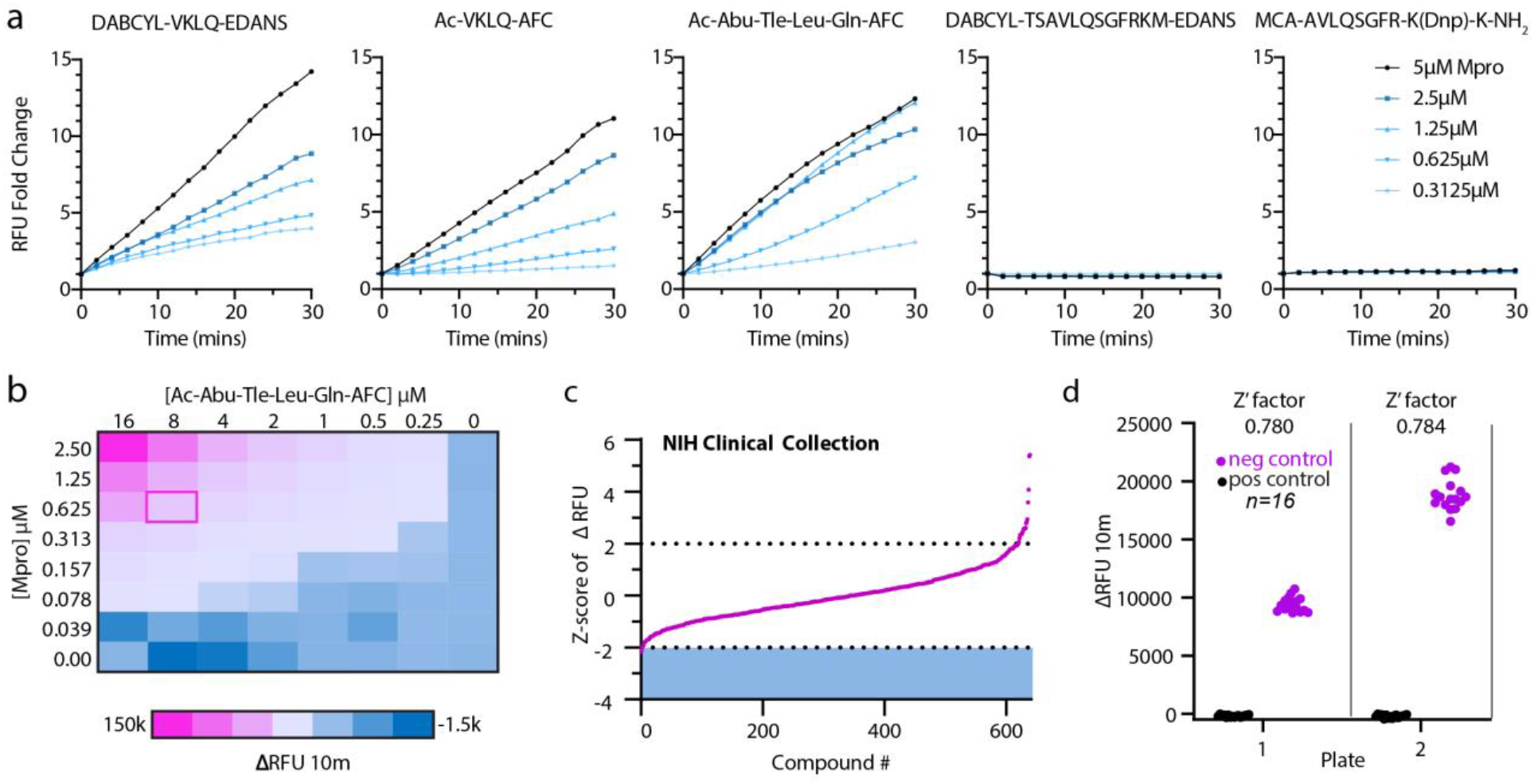
Mpro assay optimization. (a) Screen of selection of reported Mpro substrates: DABCYL-VKLQ-EDANS, Ac-VKLQ-AFC, Ac-Abu-Tle-Leu-Gln-AFC, DABCYL-TSAVLQSGFRKM-EDANS, MCA-AVLQSGFR-K(Dnp)-K-NH_2_. Fold change of increase in RFU was measured over 30 minutes holding substrate constant at 12.5µM with increasing concentrations of Mpro recombinant protein as indicated. (b) Two-dimensional titration of substrate Ac-Abu-Tle-Leu-Gln-AFC against Mpro. Concentrations are indicated for substrate along the x-axis and for Mpro along the y-axis. Heat map corresponds to the change in RFU over 10 minutes and pink outline (ΔRFU 10m=20,810) indicates chosen concentration for NIH Clinical Collection screen (0.625µM Mpro and 8µM Substrate). (c) Z-score index of NIH Clinical Collection screen (640 compounds). Hit window was considered at Z-score ≤ −2 and was calculated as the Z-score of ΔRFU at 10 minutes corresponding to the linear portion of the curve. X-axis indicates arbitrary compound number arranged by increasing Z-score. (d) Z’-factor for the two NIH Clinical Collection 384-well plates. Pink circles indicate negative control (DMSO) and black circles represent positive controls (no protein). Z’factor calculated at 0.780 and 0.784 for plates 1 and 22 respectively. Y axis represents change in RFU over 10 minutes.

**Fig 2.**
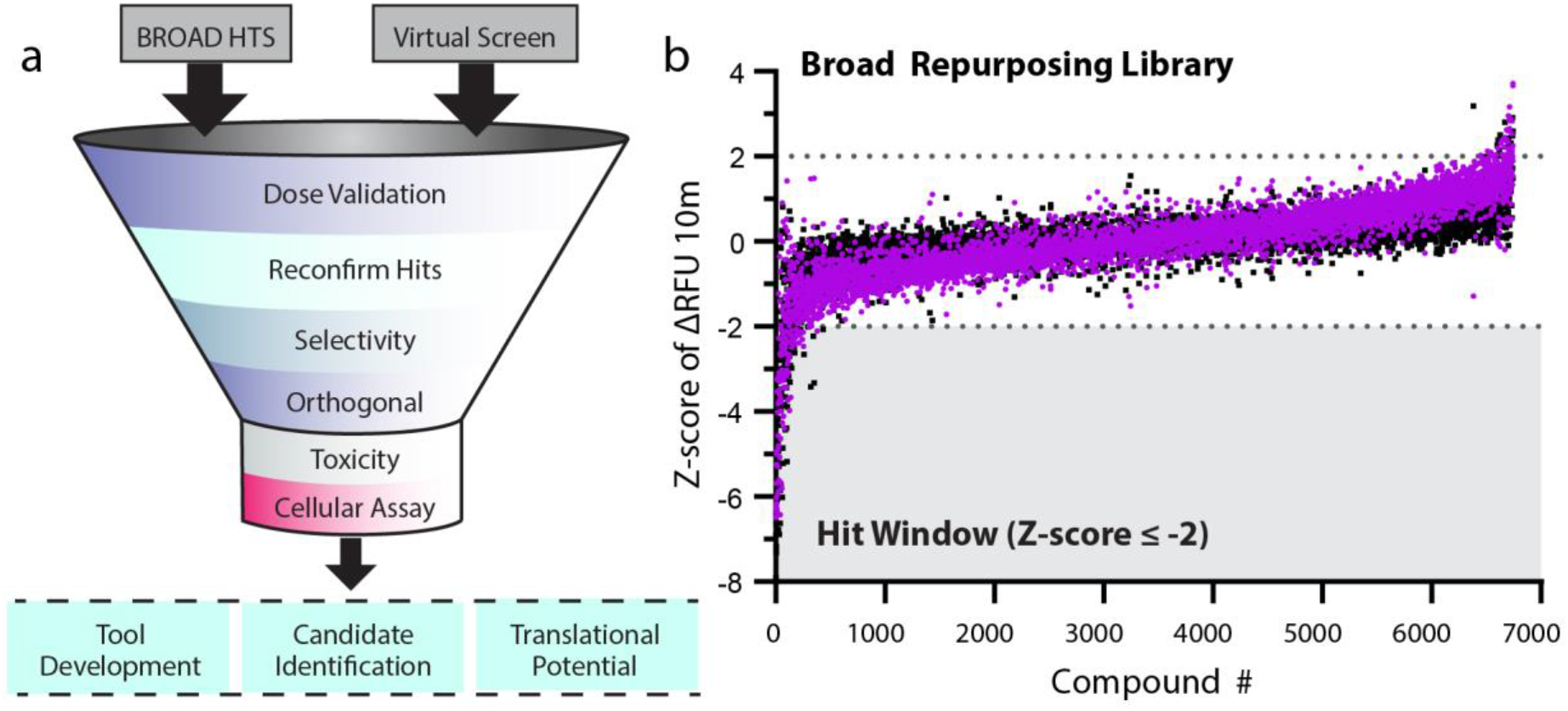
Screening pipeline and Broad library screen. (a) Schematic of screening funnel. The Broad Repurposing library was screened both empirically and virtually. Any hit from the Broad library (Z-score ≤ −2) was validated for dose-responsiveness. All suitable compounds passing this filter with satisfactory curve fitting and potency were ordered as powder and re-validated. Future efforts will test for selectivity and in orthogonal assays for suitability. Although outside the scope of this report, determination of viral anti-replicative properties as well as toxic profile at required dosage will be determined. The goal of this paradigm is to find suitable candidates for development both as tools for probing underlying mechanisms of SARS-CoV-2 as well as for translational potential. (b) Screen of the Broad Repurposing Library. Library was screened at a concentration of 50µM against both Ac-VKLQ-AFC (black) and Ac-Abu-Tle-Leu-Gln-AFC (purple). Hit window was considered for compounds falling below Z-score ≤ −2 against both substrates and consisted of 50 compounds. Compounds ordered by average Z-score.

### Drug Repurposing Strategy – screening the Broad Repurposing Library

The concept of drug repurposing is to utilize existing therapeutic drugs to treat a new disease indication. This approach is particularly relevant for COVID-19 because of the potential for an accelerated clinical impact as compared to *de novo* drug development. A systematic approach to facilitate drug repurposing has recently been described (^18^, http://www.broadinstitute.org/repurposing) and has made a large collection of drugs with previous history of use in humans available for high throughput screening. We acquired this ∼6,070 compound library as an assay ready collection in 384-well format. We conducted the library screen at 384-well density using the optimized kinetic Mpro assay described in Fig 1. Our overall repurposing strategy is described in Fig 2A. We conducted a single point screen at 50 μM compound concentration and observed ∼50 compounds with activity against SARS-CoV-2 Mpro for an overall hit rate <0.75%. These compounds were screened in parallel against the natural amino acid substrate (Ac-VKLQ-AFC) as well as a kinetically preferred substrate (Ac-Abu-Tle-Leu-Gln-AFC) (Fig 2B). Individual compounds are shown in Table 1.

**Table 1.**
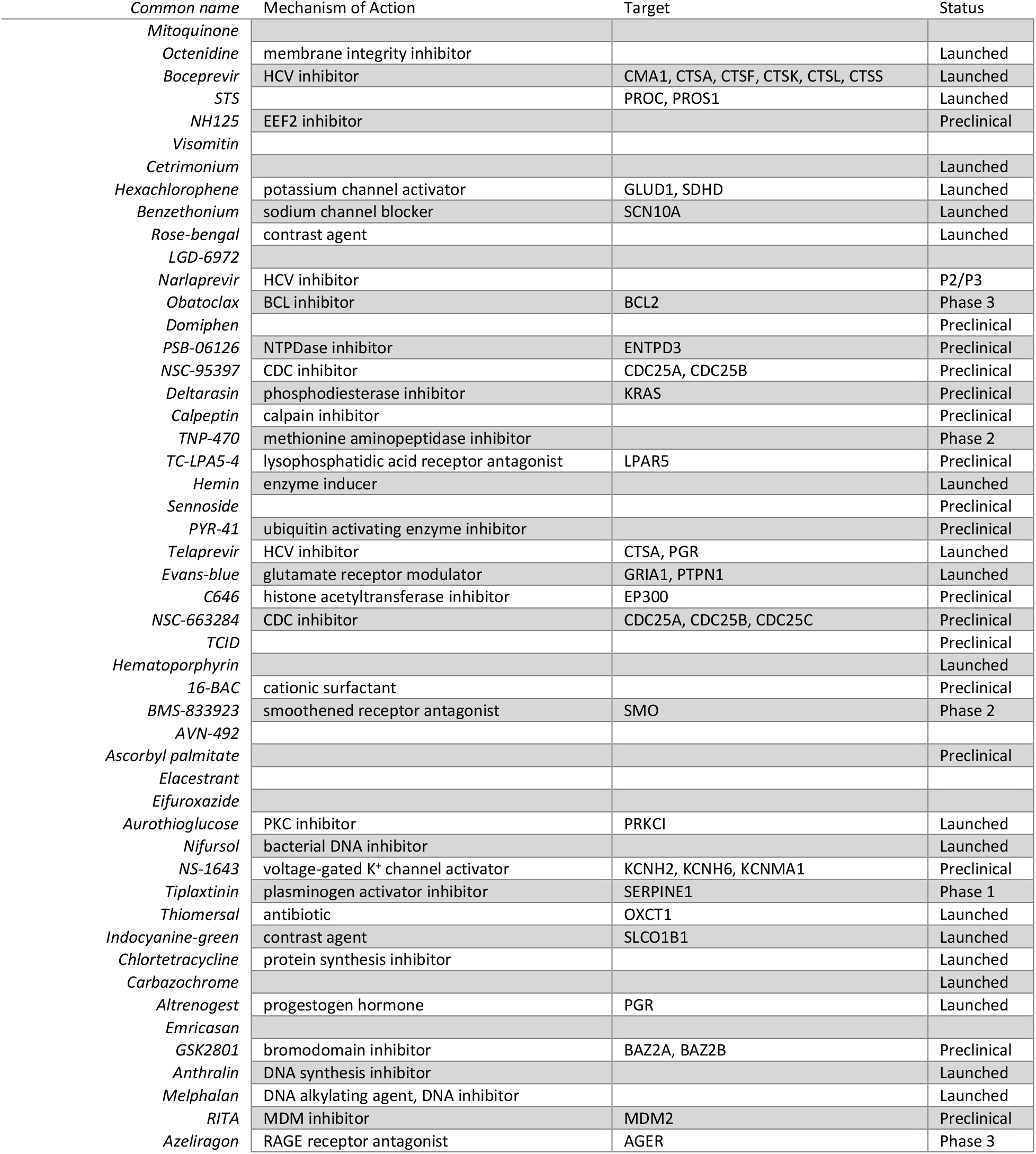
Hits identified from Broad Repurposing Library screen against SARS-CoV-2 Mpro.

### Analysis of potency

We validated the hits from the primary screen by conducting a 10-point dose-response analysis with a drug concentration range from 150 μM down to 7.6 nM (3-fold dilution series). From this dose-response analysis, IC50 values were calculated for dose-responsive hits. Several of the drugs uncovered in our screen including, Boceprevir (IC50 = 0.95 µM), Ciluprevir (IC50 = 20.77 μM), Narlaprevir (IC50 = 1.10 µM), Telaprevir (IC50 = 15.25 µM), are antiviral compounds targeting the hepatitis C NS3 protease. Boceprevir, Narlaprevir, and Telaprevir are approved drugs with a track record of safe use in human patients ^23-28^. Other relatively potent dose responsive compounds emerging from our screen include calpeptin (IC50 = 4.05 μM), aurothioglucose (IC50 = 13.32 μM), PYR-41 (IC50 = 17.38 μM), and hemin (IC50= 9.68 μM) (Fig 3).

**Fig 3.**
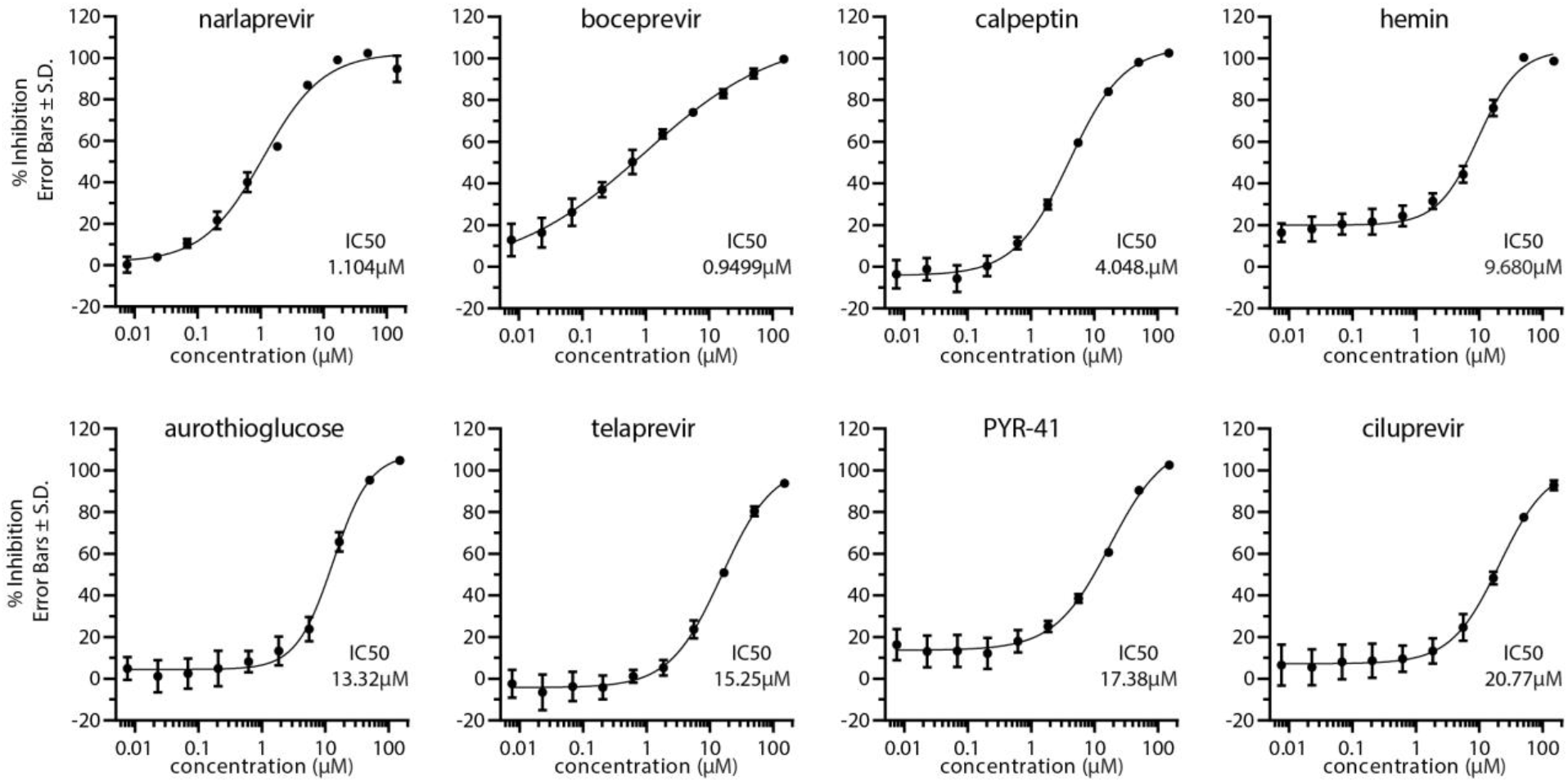
Dose-response validation of compounds against SARS-CoV-2 Mpro. Dose response curves were generated using Ac-Abu-Tle-Leu-Gln-AFC substrate. Percent inhibition of each compound was calculated at indicated 10 concentrations by comparing slope of treatment versus DMSO control. Error bars represent standard deviation and n=3 for each concentration. IC50 values are indicated and calculated by 4-parameter nonlinear regression curve fitting.

### The relative utility of *in silico* and HTS repurposing screens

The recent publication of the crystal structure for Mpro has enabled computational approaches to Mpro drug discovery ^8,9^. We leveraged the existing structural data (PBD entry 6LU7) to conduct a computational free energy calculation based *in silico* screening approach. To do this we have utilized the Schrodinger Maestro software package ^29,30^ to conduct a computational docking of all compounds in the Broad Repurposing library. Using this approach, we derived a docking score for each compound (see Table S1 for broad repurposing library with docking scores). We observe a poor correlation (Pearson r=0.02864) between Mpro docking score and Z-score in the protease inhibition assay (Fig 4A). Furthermore, top hits from the screen also exhibit a weak correlation (Pearson r=-0.1503) between compound potency and docking score (Fig 4B).

**Fig 4.**
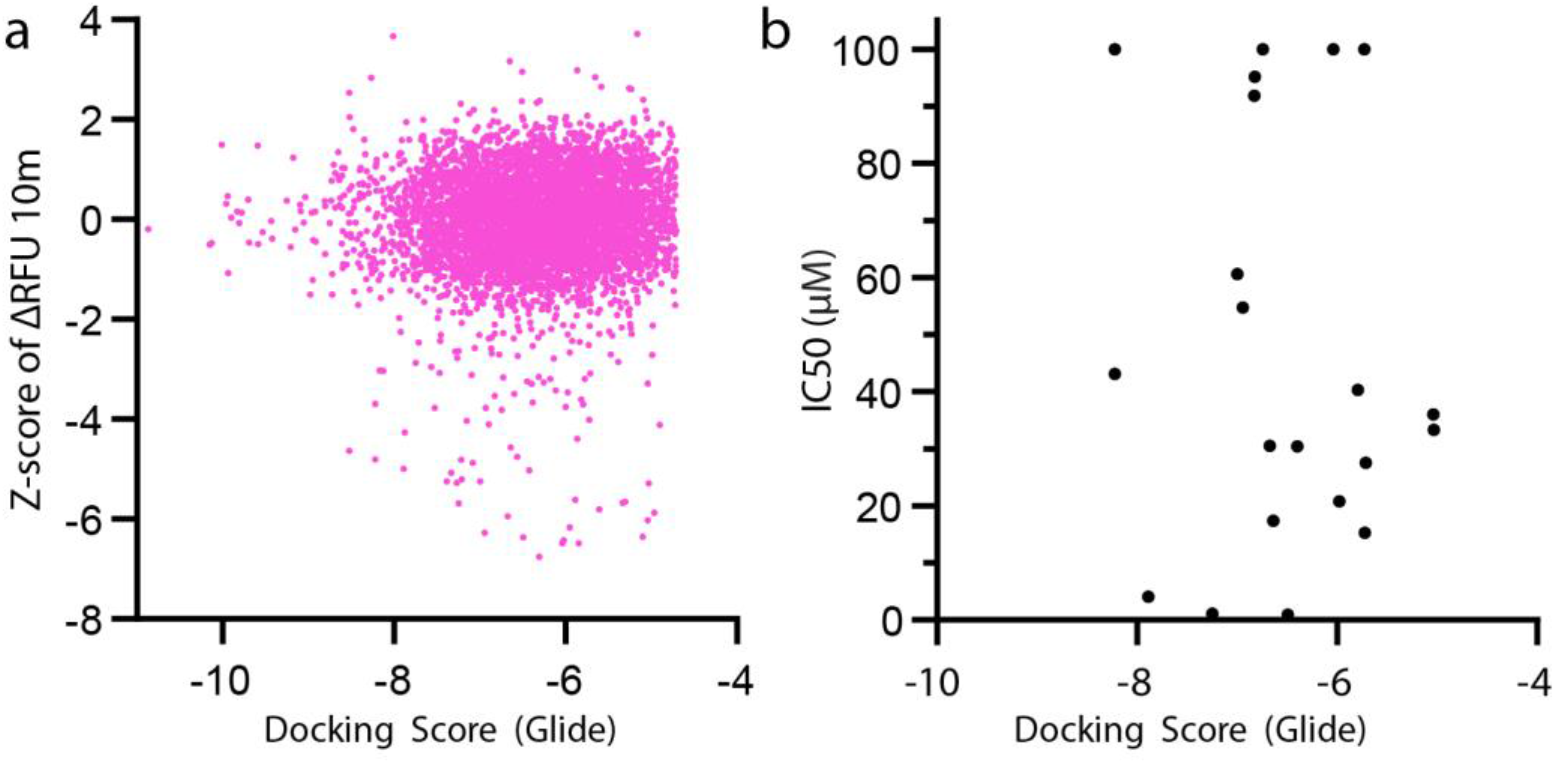
Correlation of Experimental and In Silico identification. (a) Correlation of Z-score for all Broad Repurposing library compounds and Docking Score. Pearson r = 0.02864 (95% CI 0.003478-0.05376), P value = 0.0257 and XY pairs = 6068. Linear equation Y=0.03452x+ 0.2202 as fitted by simple linear regression, slope significantly deviated from 0, P value = 0.0016. (b) Correlation of IC50 vs Docking Score. Pearson r = −0.1053 (95% CI −0.5136-0.3420), P value = 0.6498 and XY Pairs = 21. Linear equation Y=-4.188x+20.41 as fitted by simple linear regression, slope not significantly deviated from 0, P value = 0.6498.

## Discussion

### Therapeutic potential of Mpro as a target

A general discussion of drug repurposing for COVID-19 suggests viral encoded proteases may be relevant therapeutic targets for coronaviruses ^31^. Relative to most other human viruses, our understanding of the virology of SARS-CoV-2 remains incomplete. However, after decades of extensive research, we have learned a great deal about viral proteases in general and the chemical means to inhibit them from our studies of HIV-1, hepatitis C and rhinoviruses. Likewise protease inhibitors targeting the SARS protease have been investigated ^32,33^. Furthermore, previous work on SARS Mpro has demonstrated that it is a targetable enzyme worth significant translational effort ^34^.

### Perspective on hit compounds and future screening

Sequence alignment shows a high degree of homology between SARS Mpro and SARS-CoV-2 Mpro with ∼95% amino acid sequence identity. Recent studies have solved the crystal structure of SARS-CoV-2 Mpro and compared it with SARS Mpro showing that they have similar but distinct active site pockets and will require distinct dugs for potent and highly specific inhibition. This sequence and structural information has provided an opportunity to conduct *in silico* docking of known drugs to the COVID-19 virus Mpro activ site. However thus far, such analysis has not uncovered potent inhibitors of Mpro. Recent *in silico* work has suggested that protease inhibitor drugs may inhibit SARS-CoV-2 Mpro ^35,36^. However, our findings suggest that *in silico* approaches alone cannot substitute for enzyme kinetic screening evaluation of Mpro inhibitors because most identified high scoring compounds in *in silico* docking studies lack activity against Mpro in kinetic protease assays.

### Potential for Drug Repurposing and Clinical Trials

To date clinical trials for COVID-19 have not yielded potent antiviral therapies. The objective of this work is to complete a survey of approved drugs to identify therapies that can block COVID-19 viral replication by inhibiting the main viral protease. The advantage of this approach is that any approved drug identified can be advanced rapidly to clinical trials without extensive multi-year preclinical development efforts. This is also particularly germane given the limitations of animal models of COVID-19 infection and pathogenesis.

A diverse variety of initial hits were identified in our high throughput screen of the broad library. Of these, the most potent hits are all known protease inhibitors and there is strong representation from protease inhibitors developed to inhibit HCV protease NS3/4A (Boceprevir, Ciluprevir, Narlaprevir, and Telaprevir). Clearly as approved or well-developed clinical candidates, these drugs exhibit pharmacological and pharmacodynamic properties well suited to repurposing as a COVID-19 antiviral therapy. Boceprevir and Narlaprevir appear the most potent against Mpro and may be suitable for repurposing.

Previous clinical evaluation of Boceprevir (also known as Victrelis) showed it to be safe and effective for treating HCV ^37^. Boceprevir was approved as a first in class HCV NS3/4A serine protease inhibitor for treatment of chronic HCV infection. Boceprevir was FDA approved for use in the USA in 2011 and Boceprevir treatment is given as a combination therapy with interferon α2b and ribavirin. Likewise, clinical evaluation of Narlaprevir (also known as Arlansa or SCH900518) showed it to be both safe and to exhibit antiviral activity when combined with interferon α2b ^38^. Furthermore, Narlaprevir has been show effective against HCV NS3/4A mutations causing resistance to protease inhibitors ^39.^ Narlaprevir was approved for use against HCV in Russia in 2016.

Our findings demonstrate Boceprevir and Narlaprevir potency against SARS-CoV-2 Mpro in the one micromolar range. Previous work to develop Boceprevir and Narlaprevir as approved HCV therapeutics make them drug repurposing candidates worth further evaluation. Likewise, previous commercially developed NS3/4A inhibitor lead compounds may be suitable for further repurposing studies. The COVID-19 pandemic has revealed an urgent unmet medical need for potent antiviral agents for treatment of SARS-CoV-2 infection. Because antiviral therapies are frequently most effective when used in combination ^40^, it may be useful to consider combining Mpro inhibition with other antiviral strategies for treating SARS-CoV-2 infection. For instance, inhibition of the SARS-CoV-2 replicase in combination with Mpro inhibition might exhibit synergistic antiviral activity. Remdesivir, a broad-spectrum antiviral replicase inhibitor has shown efficacy against a wide variety of RNA viral replicases including SARS-CoV-2 ^41,42^. Thus, combination therapies using Boceprevir/Remdesivir or Narlaprevir/Remdesivir may yield a synergistic drug repurposing strategy for treating COVID-19. Taken together, the work presented here supports the rapid evaluation of previous HCV NS3/4A inhibitors for repurposing as a COVID-19 therapy.

## Methods

### Recombinant Protein

Recombinant Mpro was purified using constructs and methods as previously described ^8^. pGEX-6p-1 plasmid containing SARS-CoV-2 Mpro was gifted from Hilgenfeld lab at Luebeck University, Germany. Plasmid was transformed into BL21 (DE3) bacteria (NEB). A single colony was inoculated into 10Ml Terrific Broth (TB) + Carbenicillin (25ug/mL) and grown overnight to saturation. Overnight culture was transferred into 1L of TB and grown in a shaking incubator at 37°C until log phase (OD_600_∼0.7). Culture was induced with IPTG (1mM final) and kept in 37°C shaking incubator for 4 hours. Culture was spun down at 3,400rpm for 30 min at 4°C, and pellet resuspended in PBS with 10% sucrose then spun at previous conditions. PBS was aspirated and bacteria pellet was snap frozen in liquid nitrogen and stored at −70°C. Pellet was thawed and resuspended in Lysis buffer (PBS, 0.3% lysozyme, 1mM DTT, 1.5% Sarkosyl, RNAse A, and DNAse I) and sonicated for 10 seconds ON time, 20 seconds OFF time for 5 minutes of total ON time at 60% amplitude. Lysate was spun at 16,000rpm for 30 minutes at 4°C. 4mL of Ni-NTA beads and supernatant were rotated for 2 hours at room temperature. Gravity column was used for purification with His_6_-tagged Mpro binding to Ni-NTA beads (Qiagen 30210), washed with lysis buffer + 10mM imidazole and eluted with increasing concentration of imidazole (50mM, 100mM, 150mM and 200mM). The majority of Mpro eluted at between 150-200mM imidazole and was 90%+ pure by Coomassie stained gel analysis. Mpro fractions were pooled and buffer exchanged into 20mM Tris pH 7.8, 150mM NaCl, 1mM EDTA, 1mM DTT, and snap frozen in liquid nitrogen and stored at −70°C. Yield and purity were assessed via BCA (ThermoFisher 23225) and Coomassie-stained SDS-PAGE.

### Fluorescent Mpro Protease Assay

Fluorescent Mpro peptides were synthesized by Anaspec (www.anaspec.com) at 90% purity and frozen in 1 mg aliquots. Stock concentration of substrates were made by reconstituting powder in 100uL DMSO (10mg/ml) and storing at −70°C. Optimal Mpro substrates were previously determined to be Ac-Val-Lys-Leu-Gln-AFC for physiological substrates and Ac-Abu-Tle-Leu-Gln-AFC, a noncanonical amino acid substrate ^20^. Ac-Val-Lys-Leu-Gln-AFC and Ac-Abu-Tle-Leu-Gln-AFC fluorogenic substrates were monitored at 380/20 nm excitation and 500/20 nm emission wavelengths. FRET-based substrates Dabcyl-Val-Lys-Leu-Gln-EDANS was measured at 336/20 nm excitation and 490/20 nm emission wavelengths and MCA-Ala-Val-Lys-Gln-Ser-Gly-Phe-Lys-DNP-Lys was monitored at 325/20 nm excitation and 392/20 nm emission wavelengths. We used 20mM Tris pH 7.8, 150mM NaCl, 1mM EDTA, 1mM DTT, 0.05% Triton X-100 as the assay buffer. Assay conditions were at room temperature (25°C) for all assays.

### 2D Titration main screen optimization

2D titration for determining the main screen ratios was done in 96 well black opaque plates (Corning 3686 NBS). The top concentration of Mpro was 2.5µM and serial diluted to 0.0395µM along the Y-axis of the plate. The top concentration of substrate was 16µM and serial diluted along the X-axis of the plate. Fluorescence was monitored at 380/20 nm excitation and 500/20 nm emission wavelengths.

### Broad Repurposing Library

The Broad Repurposing Library was ordered and plated into black opaque 384-well plates (Greiner 781209) at 100nL of 10mM (slight variations depending on compound) compound in DMSO. 10uL of diluted Mpro (625nM final concentration in reaction buffer detailed above) was added with a MultiFlo FX liquid dispenser using a 5µL cassette. Compounds were incubated with Mpro for 10 minutes at RT after which 10uL of substrate (8uM final concentration of either Ac-VKLQ-AFC or Ac-Abu-Tle-Leu-Gln-AFC) was dispensed into the plate and read using a Cytation 5 multi-mode reader immediately at 380/20 nm excitation and 500/20 nm emission wavelengths every 5 minutes for 30 minutes. Data was analyzed using Biotek Gen5 software, Microsoft Excel, and GraphPad Prism 8.

### Dose validation assays

Hit compounds were ordered from the Broad Institute pre-plated in 384-well format (Greiner 781209) as 10-point serial dilutions (3-fold) at 300nL per well. Mpro (80nM final concentration) and substrate (Ac-Abu-Tle-Leu-Gln-AFC at 32µM final concentration) were dispensed in the same manner described above. Inhibition was calculated as 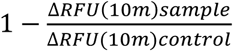 at each concentration and data fitted to 4-parameter nonlinear regression model using GraphPad Prism 8.

### *In Silico* Docking of the Broad Repurposing Library with Mpro

We utilized the Schrodinger Maestro software package ^29,30^ to conduct a computational docking of all compounds in the Broad Repurposing library. In this approach we generated a receptor grid model of the Mpro active site and serially docked each compound in the Broad Repurposing library with the active site model using the physics based Glide algorithm ^43,44^. We chose the Glide algorithm over the many competing options because of its superior performance in head to head comparisons of algorithms ^43^.

### Statistical analyses, IC50 Calculation, Selectivity Calculation, and figures

Graphs were generated using GraphPad Prism 8. IC50 calculations were performed using GraphPad Prism 8 curve fitting using 4-parameter non-linear regression.

## Supporting information

Screen Summary Data

## Author Contributions

J.D.B. and B.C.K. designed research; J.D.B., R.L.U. performed research; J.C.K performed *in silico* docking. J.D.B. and B.C.K. analyzed data; B.CK. & J.D.B. wrote the first draft of the manuscript. All authors reviewed, edited, and approved the final version of the manuscript.

## Acknowledgements

We thank Timothy Strovas, Aleen Saxton, Caleb A. Baker, Misa Baum, and Jeanna Wheeler for sharing their expertise. We thank the Center for the Development of Therapeutics and Repurposing Hub at the Broad for providing the compounds library (https://www.broadinstitute.org/repurposing). We thank James Meabon for assistance with acquiring and maintaining critical HTS infrastructure.

## Data Sharing

Raw data from the primary screen of the Broad library is available online in the Sage Biosystems data repository, synapse.org (see: https://www.synapse.org/#!Synapse:syn22275070/files/).

